# Fusion of single-cell transcriptome and DNA-binding data, for genomic network inference in cortical development

**DOI:** 10.1101/2021.05.18.444638

**Authors:** Thomas Bartlett

**Affiliations:** University College London

**Keywords:** Gene regulatory networks, single-cell RNA-seq, cortical development, computational biology

## Abstract

**Background:** Network models are well-established as very useful computational-statistical tools in cell biology. However, a genomic network model based only on gene expression data can, by definition, only infer gene co-expression networks. Hence, in order to infer gene regulatory patterns, it is necessary to also include data related to binding of regulatory factors to DNA.

**Results:** We propose a new dynamic genomic network model, for inferring patterns of genomic regulatory influence in dynamic processes such as development. Our model fuses experiment-specific gene expression data with publicly available DNA-binding data. The method we propose is computationally efficient, and can be applied to genome-wide data with tens of thousands of transcripts. Thus, our method is well suited for use as an exploratory tool for genome-wide data. We apply our method to data from human fetal cortical development, and our findings confirm genomic regulatory patterns which are recognised as being fundamental to neuronal development.

**Conclusions:** Our method provides a mathematical/computational toolbox which, when coupled with targeted experiments, will reveal and confirm important new functional genomic regulatory processes in mammalian development.

## Background

Network models have become very popular in cell biology in recent years, proving their usefulness in many contexts. Example applications include gene regulatory, co-expression and protein signalling networks. Most applications in cell biology continue to use static network models, including in the context of single-cell RNA-seq data [1, 2]. However, processes such as development are inherently dynamic, and hence for such applications, a time-varying genomic network model would be more appropriate. Better inference of time-varying genomic networks will allow regulatory patterns to be inferred which better characterise dynamic biological processes such as development.

Developmental processes are characterised by transient expression of certain key genes at specific times. Morphogen gradients set up at particular developmental stages provide specific information about location of cells, leading to appropriate patterns of gene expression in those cells. These gene expression patterns define cellular lineages, which can then be locked in place by persistent expression of, for example, homeobox and bHLH genes [3]. In the neural lineage, particular subtypes of fully differentiated neurons and glial cells are arrived at as a result of sequential expression of such positional markers and fate-determining genes [4, 5, 6, 7].

In the cortex of mammalian embryos, early in neural development morphogen gradients driven by WNT and SHH signalling are set up defining location in the cortex. For example, SP8 and COUP-TFI are expressed most strongly at either end of a rostrodorsal to caudoventral gradient, and PAX6 and EMX2 expressed most strongly at either end of a rostroventral to caudodorsal gradient. The positional information from these morphogen gradients leads to specific developmental trajectories being followed in human embryos, involving the expression of numerous other master regulators and positional markers such as NKX2-1, SOX6, COUP-TFII, GSX2, DLX1/2, and OLIG2 [8]. For example, later in neural development, these trajectories can lead (depending on location) to the sequential expression RELN, TBR1, CTIP2, CUX1 and SATB2 [4], which determine the specification of excitatory neuron subtypes. For a more detailed background on the molecular mechanisms of neuronal specification, a thorough review is provided by Guillemot and colleagues [9].

Cells can be characterised in terms of their progression through a dynamic process such as neural development according to their gene expression patterns. In this context, cells with similar gene expression patterns are characterised as being at a similar point along the developmental process, or trajectory. The notion of ‘developmental time’ of cells can be used to quantify the progression of those cells through the developmental process, or trajectory. Hence, developmental time can be characterised in terms of the gene expression patterns of the cells. It is also recognised that genomic network inference in single-cell data should be carried out on cells of specific types [10]. A time-varying network model allows the dynamic genomic network structure to be inferred from relatively homogenous groups of cells, with each such group of cells corresponding to a different developmental time-point. In this model, each such group of cells represents a different time-step along the developmental process, or trajectory.

Any genomic network model based only on gene expression data can by definition only infer gene co-expression networks. In order to infer gene regulatory patterns, it is necessary to also include data which relates to the physical binding of the products of some genes to DNA of other genes. Previous work has been successful at inferring these more complicated genomic network structures, by incorporating data of several different modalities [11, 12]. However, those models are very computationally intensive, and are appropriate only for small networks involving the influence of few tens of gene regulators (such as transcription factors) on a few hundreds of genes. On the other hand, the method which we propose here is able to infer genomic regulatory patterns from genome-wide data, based on the expression of tens of thousands of genes and / or transcripts. However, we also note that to confirm any novel findings of marker genes or transcription factors important in neurogenesis, any analysis using this method will need to be coupled with experimental verification. This would require a combined dry/wet lab setting, which is beyond the scope of this investigation.

In this work, we propose a method to infer dynamic genomic network structure, fusing single-cell RNA-seq data from specific experiments (i.e., gene expression data) with publicly-available DNase-seq data (i.e., DNA-binding data). This paper is organised as follows. In Section 1, we define our model, and describe our inference method. In Section 2, we present the results of applying our model/method to data from human fetal cortical development. Then in Section 3, we discuss our findings and their wider implications.

## 1 Methods

### 1.1 Model overview

We infer genomic network structure by fitting a sparse linear model locally around each ‘target gene’. This sparse linear model has the log gene expression for target gene *i* at time *t* as the response, and the log gene expression for all genes *j* ≠ *i* genome-wide at time *t* as potential predictors; variables are standardised before model fitting. From this genome-wide choice of potential predictor genes, the sparse model fit chooses a small set of predictor genes which together are able to predict the expression level of the target gene *i*. This chosen set of genes are then used to infer the local network structure around the target gene *i*. To infer the global network structure, we infer the local network structure around each target gene *i* in turn.

As well as gene expression data, we also use DNA-binding data to inform the sparse model fits. We use this DNA-binding data to reduce the sparsity of the model fit for predictor genes for which there is evidence of a physical DNA interaction between the gene-product of predictor gene *j* with the DNA of target gene *i*. This means that the sparse model fit is more likely to infer genomic network interactions between predictor genes and a target gene whenever there is evidence of a physical interaction between the gene product of the predictor genes and the DNA of the target gene.

### 1.2 Time-varying network model

Following earlier work [13], denoting log(gene expression+1) at time *t* as *y_t_* for the target gene *i* and **x**_*t*_ for the *p* − 1 other genes, the model is defined as:

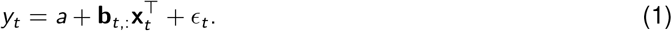

The time-varying coefficient vector **b**_*t*,:_ encodes the time-varying local network structure at time *t* around the target gene. If there is a non-zero element of this vector at *b_t_*_,*j*_ (after thresholding to remove trivially small values), then a network edge is inferred between genes *j* and *i*. The row-vector **b**_*t*_, : is a row of the matrix **b**, and hence **b** encodes the time-varying local network structure around the target gene *i* for all times *t* ∈ {1,…, *T*}.

Assuming local decomposability of the global network structure as in [14] allows the local network structure to be inferred separately around each target gene *i*. Also assuming Gaussian distributed errors with constant variance leads to the log-likelihood

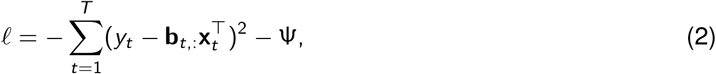

where the subtraction of the ψ term leads to ‘regularisation’, or ‘penalisation’ (of the model likelihood). A special case of the model, which assumes no DNA-binding data is available to adjust local sparsity, defines ψ as

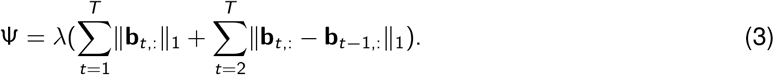

The first term of ψ, i.e., 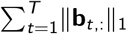, encourages choosing a smaller number of regulator genes, minimising the number of non-zero entries in **b**_*t*,:_, which is referred to as ‘sparsity within time’ [13]. The second term of ψ, i.e., 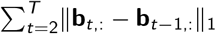, encourages smooth time-variation of **b**_*t*,:_, which is referred to as ‘sparsity across time’ [13].

The type of likelihood penalisation specified by ψ (Equation (3)) falls within the generalised lasso framework [15], meaning that ψ can be written as

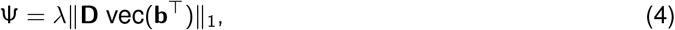

where vec(·) is the vectorisation operator (which vectorises the (*p* − 1) × *T* matrix **b**^*T*^ to a (*p* − 1)*T* × 1 vector), and *λ* controls how sparse the model is. The penalty matrix **D** ∈ ℝ^*m×*(*p−*1)*T*^ controls which elements of **b** are sparse, as well as controlling which differences between elements of **b** are sparse. Each row of **D** defines a different component of the sparsity. If exactly one element of a row of **D** is non-zero, then this leads to a contribution to the sparsity within time, i.e., sparsity for the variable and time-point at the corresponding location in vec(**b**^*T*^). If exactly two elements of a row of **D** are non-zero, are of equal magnitude but opposite sign, and correspond to locations in vec(**b**^*T*^) for the same variable at adjacent time-points, then this leads to a contribution to the sparsity across time. These are the only scenarios for this model in which any element of **D** is non-zero.

To achieve sparsity within time, there must be a separate row in **D** for each gene *j* for each time-point, with one non-zero element in each of these rows. The level of sparsity can be varied for each gene *j* by varying the magnitude of these non-zero elements, as long as the magnitude of the non-zero elements is the same for all *T* rows of **D** which correspond to gene *j*. Similarly, to achieve sparsity across time, there must be a separate row in **D** for each gene *j* for each pair of adjacent time-points *t* − 1 and *t*, with *t* ∈ {2,…, *T*}. Each of these rows must have exactly two non-zero elements with the same magnitude and opposite signs, at locations corresponding to times *t* − 1 and *t* for gene *j*. Again, the level of sparsity can be varied for each gene *j* by varying the magnitude of these non-zero elements, as long as the magnitude of the non-zero elements is the same for all *T* − 1 rows of **D** which correspond to gene *j*. In practice, we set the magnitude of the non-zero elements to be the same for all 2*T* − 1 rows of **D** which correspond to gene *j* - this covers both sparsity within and across time.

### 1.3 Using DNA-binding data to adjust local sparsity

To set the magnitude of the non-zero elements of the rows of **D** which correspond to gene *j*, we use the evidence available in DNA-binding data for any interaction of the protein-product of gene *j* with the promoter DNA of the target gene *i*. We set these magnitudes from model probabilities which quantify the evidence in the DNA-binding data for this protein-DNA interaction. These fitted model probabilities can come from any model, meaning that the framework presented here is independent of the model of the DNA-binding data which is used. Other authors have previously used such a notion of model independence for data-fusion in genomics [12].

The magnitude of the non-zero elements of **D** defaults to 1 whenever there is no evidence for the binding of the protein-product of gene *j* to the promoter DNA of the target gene *i*. This is typically the case when, for example, gene *j* does not code for a transcription factor. When there is evidence of binding of the protein-product of gene *j* to the promoter DNA of the target gene *i*, the magnitude of the corresponding elements of **D** is decreased below 1. The amount by which this magnitude is decreased varies according to the strength of evidence of DNA binding, with a minimum magnitude of 1*/η* (for *η >* 1) when the binding evidence is strongest. This means that the sparsity is decreased for gene *j* according to the strength of evidence of an interaction between the protein product of gene *j* with the promoter (or other regulatory) DNA of the target gene *i*.

We assume that the model of the DNA-binding data gives binding probabilities *p_ji_* ∈ [0, 1] for the interaction of the protein-product of gene *j* with the promoter DNA of target gene *i*. Then, we want *p_ji_* = 1 and *p_ji_* = 0 to correspond to magnitudes of 1*/η* and 1 respectively, for the non-zero elements in the rows of **D** which correspond to gene *j*. For 0 *< p_ji_* < 1, we want these magnitudes to scale proportionally to log(*p_ji_*). The intuition behind this proportionality of scaling is simply that by referring to the log-likelihood in Equation (2), we can observe that the sparsity term ψ is on the scale of log-probability. Hence we want any additive components of ψ (as in Equations (3-4)) to scale proportionally with log-probabilities.

To achieve this scaling between 1*/η* and 1 proportionally to log(*p_ji_*), we set the magnitude of the non-zero elements of the rows of **D** which correspond to gene *j* as

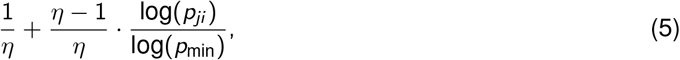

for *j* ∈ {*j*: *p_ji_ ≥ p*_min_,}, where *p*_min_ *>* 0 is the minimum model probability of interest. We set the magnitude of the non-zero elements of **D** to be 1 otherwise. Thus, *η* represents the factor by which *λ* is scaled down from gene *j* to gene *j*′, when *p_ji_* ≤ *p*_min_ and *p_j′i_* = 1. Like *λ*, the scaling factor *η >* 1 is set by the user. We have found that *η* = 4 works well in practice, and we previously found that *λ* = 20 is optimal [13].

### 1.4 Modelling co-regulation to achieve consistency in sparse model fits

The lasso is a linear model with *L*_1_ penalisation, such as in Equation (3) where ψ is comprised of ||·||_1_ terms. It is well known that lasso models may lead to inconsistent results, if the same model is fit several times to the same data under slight perturbations of the set of variables available to the model [16]. This inconsistency happens because there may be many subsets of the available variables which produce model fits which are virtually as good. A novel way to overcome this inconsistency in the context of genomic network inference is to look for genes which are consistently inferred as regulators of many genes in a particular gene-set.

In the time-varying genomic network model described here, a gene is inferred as a regulator of a target gene *i* at time *t* if it is included in the genes *j* which are selected as predictors of target gene *i* according to the non-zero model coefficients |*b_t_*_,*j*_| > 0. Genes *j* which are inferred as predictors of target gene *i* are then inferred as being connected to this target gene in the local network structure around this target gene. If a particular gene *j* is inferred in this way as being connected to many such target genes *i*, where those target genes make up a particular gene-set, then consistency is demonstrated in the regulation of the target genes *i* of this gene set by the gene *j*. Such a gene-set could correspond to, for example, the marker genes known to specify particular cell-types.

## 2 Results

### 2.1 Inference of developmental pseudo-time

The main neuro-developmental data-set analysed here [17] consists of transcriptome measurements from single cells from developing fetal brains. Some of these cells are neural stem-cells, some are fully differentiated neurons and other cell types, and there is a whole spectrum of cells in between. No information is available for each cell other than its gene expression measurements. We inferred a time-ordering of all the cells before model fitting: this time-ordering gives the relative position on a ‘developmental trajectory’ which goes from neural stem cell (inferred time 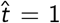) to fully differentiated cell type 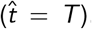. These developmental trajectories also branch with lineage, as cells are specified and differentiate into various different cell types. A time ordering inferred like this is often referred to as ‘pseudo-time’. Several methods have been published previously, to carry out this pseudo-time inference [18, 19, 20, 21]. We have followed a common theme amongst these methods, summarised as follows: (1) Dimensionality reduction e.g., by *t*-SNE (*t*-distributed stochastic neighbour embedding) [22]. (2) Trajectory and branch inference (often after some clustering, to assign cells of the same phenotype to the same pseudo-time point). (3) Biological inference (using prior knowledge to relate trajectory extrema to known cell-types).

Figure 1 shows a developmental pseudo-time ordering according to the strategy described above, based on the gene-expression measurements for the 2136 cells of the main neuro-developmental data-set analysed here [17]. The trajectories in Figure 1 are inferred from a minimum spanning tree, where the root represents the location of the stem cells and the leaves represent the fully differentiated cell types. This minimum spanning tree is fitted to the medioids of clusters obtained by the ‘partition around medioids (PAM)’ method, resulting in *T* = 8. The visualisation is via a *t*-SNE projection into three dimensions. Figure 1 also shows the cells coloured according to mean expression of marker genes identified for different cell-types (Table S1). N.B., these marker genes (provided by domain-experts) were not used in any way to infer the pseudo-time ordering or the developmental trajectories, other than to determine which end of the trajectory is the stem cell. Hence these colourings blindly verify the appropriateness of the inferred developmental pseudo-time orderings.

**Figure 1:**
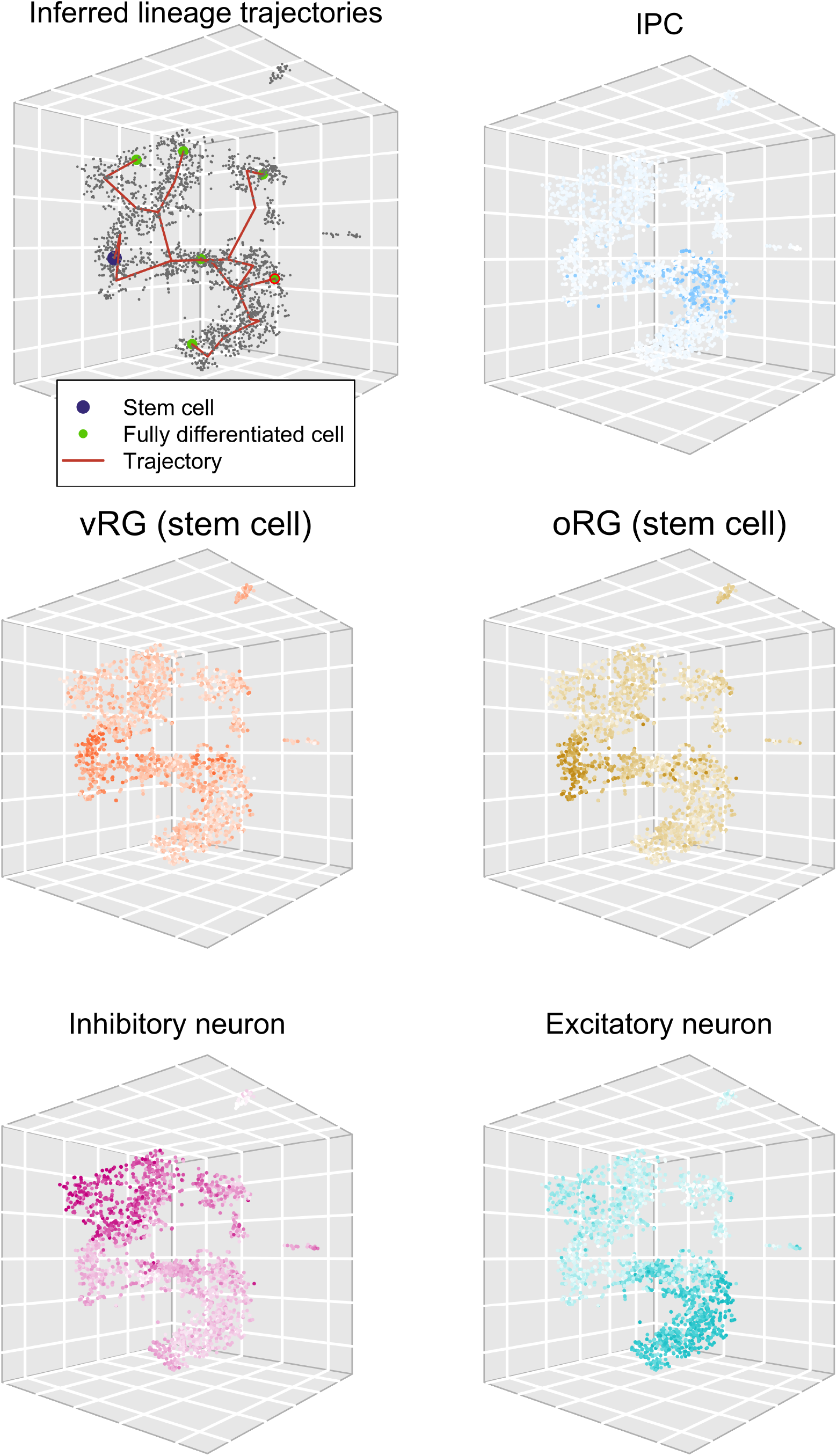
Inferred lineage trajectories, together with cellular identities for verification of trajectories. The six detected terminally-differentiated neuronal subpopulations are shown in green, with a red outline for the one analysed further in Section 2.3. vRG and oRG refer to ‘ventricular radial glia’ and ‘outer radial glia’ respectively.

### 2.2 Inference of promoter-DNA binding

In order to fit the time-varying genomic network model of Section 1, we need to obtain probabilities quantifying the evidence for the interaction of the protein-product of gene *j* with the promoter DNA of target gene *i*. We denote these probabilities *p_ji_* ∈ [0, 1], and we estimate them as the posterior probabilities of this protein-DNA interaction using the *CENTIPEDE* model [23]. We use the *CEN-TIPEDE* model with 14 DNAse-seq data-sets from human fetal brain tissue, downloaded from *ENCODE* (*www.encodeproject.org*). This provides us with the posterior probabilities of each of 415 transcription factors *j* interacting with the DNA of 13907 target genes *i*. To fit the *CENTIPEDE* model, we specify that binding should be within 5000 base-pairs (5kbp) upstream of the transcriptional start site, with a 90% minimum probability weight matrix match score. These probability weight matrices were downloaded from the *JASPAR* database (*jaspar.genereg.net*).

### 2.3 Inferring local structure with the proposed time-varying network model

The developmental trajectories inferred according to Section 2.1 were used to obtain a time-stamp for each cell. Using these time-stamps, together with with the promoter-DNA binding inference of Section 2.2, the time-varying genomic network model of Section 1 was fit to the main neuro-developmental single-cell RNA-seq data-set [17]. This model fit infers the local network structure around a target gene, by choosing which genes best predict the expression of the target gene. The model makes this choice from all other 10774 genes, genome-wide, which are present in both the main neuro-developmental data-set and the DNA-binding data-set. This model fitting procedure takes 35 minutes on one processor core (MacBook Pro, 2019, 2.6 GHz). The model can be fitted to each target gene in turn from a panel of genes of interest, or genome-wide, to give the local network structure around each target gene. We note that this procedure can easily be run in parallel on multiple cores for a large panel of target genes of interest. Figure 2 shows the inferred time-varying network structure, which resulted from fitting a single model around the target-gene SATB2.

**Figure 2:**
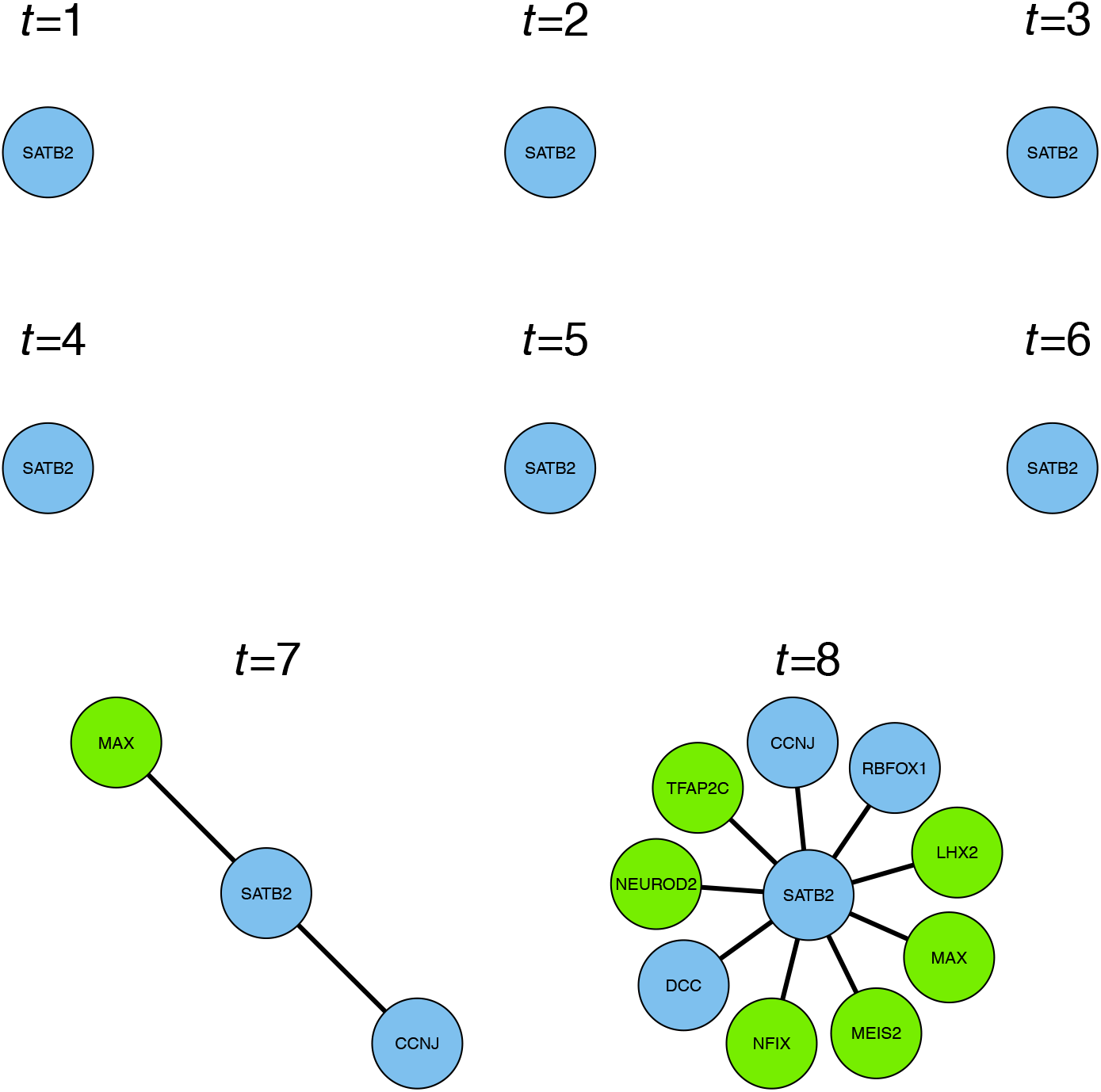
Time-varying local network structure inferred around the target gene SATB2 with the model of Section 1. Genes with transcription-factor binding data are shown in green, and other genes are shown in blue.

The gene SATB2 is well known for defining the identity of certain types of neuron [24, 4]. As would be expected for such a gene, network edges (representing direct or indirect genomic regulatory effects) appear at later times (Figure 2), as the cells take on their neuronal identities. These network edges connect SATB2 to several genes, which are all relevant to neuronal development, as follows. At the final time-point (Figure 2), NEUROD2 is recognised as the prominent neurogenesis gene ‘neurogenic differentiation factor 2’ [25]. The transcription factor NFIX is essential for neural development [26], and has a role in both embryonic and adult neurogenesis [27]. The transcription factor TFAP2C is part of the core cortical development programme [28]. The transcription factor LHX2 is known to promote neuronal as opposed to glial fate [29], as well as regulating the timing of cortical neurogenesis [30]. MEIS2 is known as a co-factor of the ventral neural fate marker PAX6 in neurogenesis [31] (PAX6 being one of the most important genes in neurogenesis [32]). The gene RBFOX1 is well known for regulating alternative splicing in neuronal development [33, 34]. The gene DCC encodes an axon guidance receptor which is important for the migration of developing neurons [35]. Then at the penultimate time-point (when the cells may still be proliferating), the transcription factor MAX is recognised as being involved with cellular proliferation [36]. Also, the gene CCNJ is a cyclin, and thus it is involved in cellular proliferation via its role in the cell-cycle.

### 2.4 Inferring network structure using the thresholded correlation matrix - a comparison

A very popular way to infer gene co-expression network structure is by thresholding the gene expression correlation matrix, e.g. at |*ρ*| ≥ 0.5, where *ρ* is the Pearson or Spearman correlation coefficient. This method of inferring genomic networks is often used in the most high-profile studies [37]. We compare this thresholded correlation matrix method of inferring genomic networks, with the proposed time-varying genomic network inference method. To make this comparison, we have inferred the local network structure around the target gene SATB2 by thresholding the correlation matrix of the cells assigned to the final time-point in the developmental trajectory (i.e., *T* = 8 in Figure 2). If we choose to threshold at |*ρ*| ≥ 0.5, we do not find any edges in the local network structure around SATB2 from this method. So instead, we threshold at |*ρ*| ≥ 0.4. The local network structure inferred in this way is shown in Figure 3. Notably, there are no well-known neuro-developmental transcription factors amongst the genes in this local network structure. Many (but not all) of the genes in this local network structure have been associated previously with neural development and brain function, although the functional role is often less clear than those shown in Figure 2. The role of the genes of Figure 3 is summarised as follows. DOCK7 is thought to help regulate radial glial proliferation [38] (radial glia are an important type of cortical stem cell). MSI1 is known to be expressed in the sub-ventricular zone of neural stem cells [39]. RPL35, SYT11 and DMTF1 have been associated in a previous bioinformatic analysis with neurogenesis [40]. CNTLN has been correlated with RB-related protection from cell division during neurogenesis [41]. FGFBP3 has been previously associated with radial precursor cells [28]. TOP2A has previously been found to be expressed in the fetal telencephalon [42]. CELF5 is known to be expressed in the brain [43]. PCDH9 is a procadherin which may be expressed in the embryonic central nervous system [44]. PSIP1 has previously been associated with hereditary hearing loss [45]. Also, expression of PHLDA1 has been correlated with intractable epilepsy [46]. We also note that the inference of these gene co-expression network patterns is static, i.e., they do not vary with time; in other words the co-expression network is constrained so that no variation with time is possible. By using a softer constraint over time, smooth variation with time is possible.

**Figure 3:**
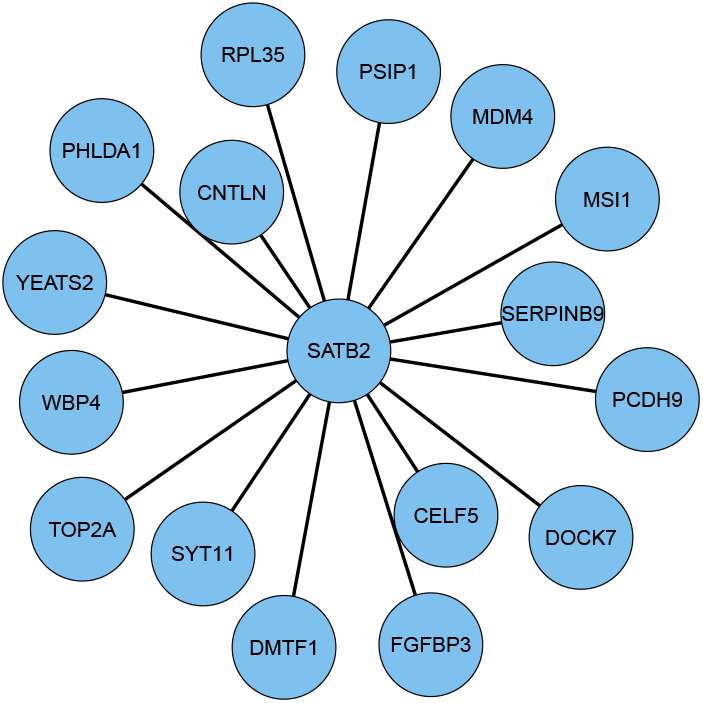
Local network structure inferred around the target gene SATB2 from the correlation matrix of cells assigned to the final time-point.

### 2.5 Consistent regulation across excitatory neuron markers

The model used to infer these results (Equations (2)-(3)) is based on the lasso [47], which selects a smaller set of predictor variables (i.e., genes in this case) for the fitted model, from the full set of variables available (i.e., genome-wide in this case). A well known property of lasso-based models is that there may be several possible sets of predictor variables which can be chosen by the model, each of which fit the data virtually equally well. Recent work has tried to overcome this issue, by looking for consistency amongst the chosen sets of predictor variables across many model fits [16]. In that work, the authors fit the model to several slight variations of the sets of predictor variables which are available to choose from, to find predictor variables which are chosen consistently across these model fits. An alternative approach which we use here is to fit the model to several different target genes, looking for predictor genes which are chosen consistently across these model fits. Importantly, these target genes are all taken from a particular gene-set of interest. This means that consistency across several model fits informs us about genomic regulatory processes which are fundamental to that gene-set of interest. Such a gene-set, comprising a highly curated selection of genes which is known to be important for excitatory neuronal identity, is given in Table S2 in the Appendix. Figure 4 shows the numbers of genes from this excitatory neuron gene-set which are found to be regulated by different transcription factors at the final time-point in the inferred dynamic network structure (the list of genes inferred as regulated by each TF is shown within the relevant bar of Figure 4).

**Figure 4:**
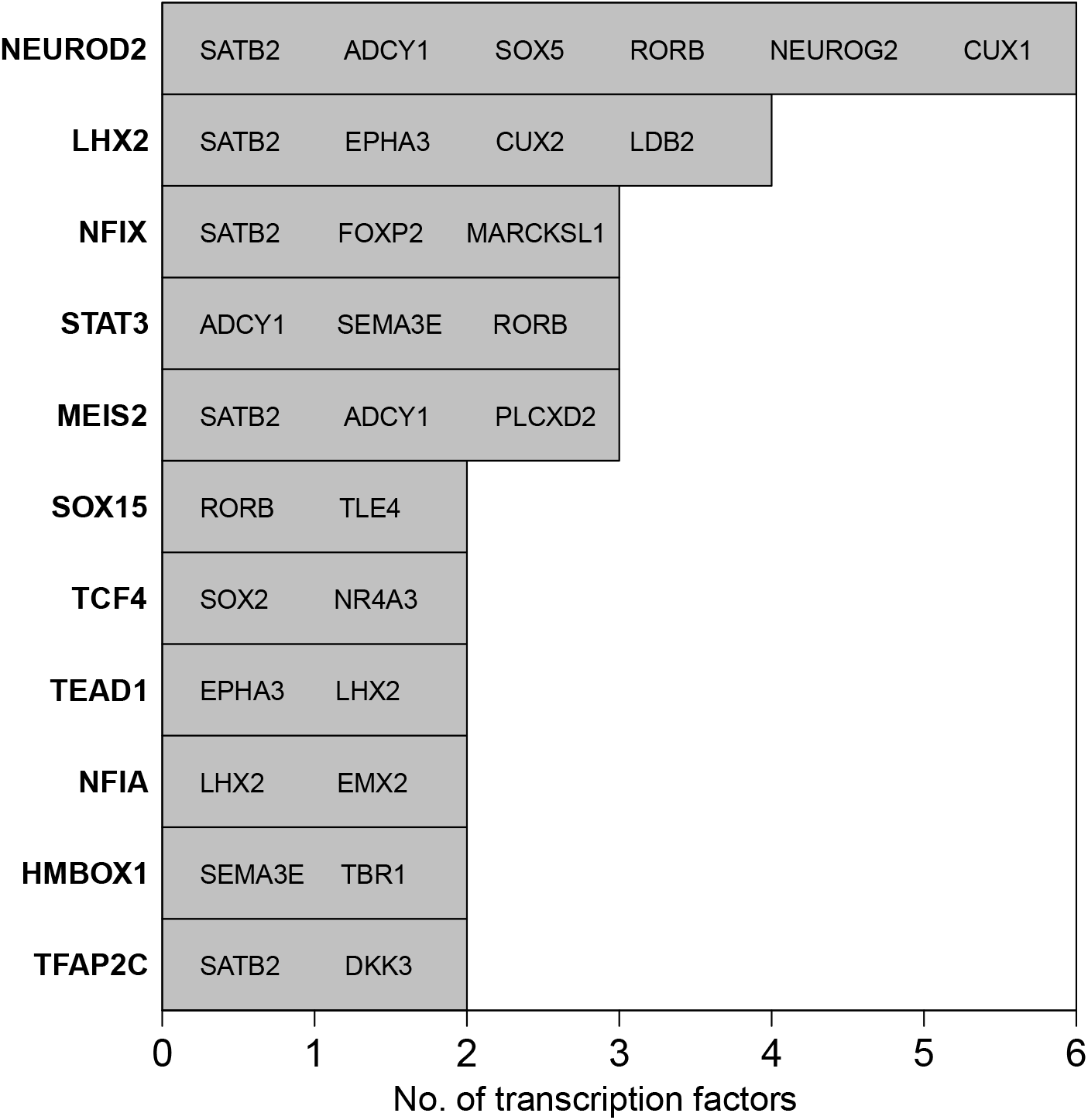
Numbers of excitatory neuron genes from the selection in Table S2, regulated by different transcription-factors.

The transcription factors which appear in Figure 4 are found to consistently regulate the genes of the excitatory neuron gene-set (Table S2 in the Appendix) and are known as being important for neural development, as follows. NEUROD2, LHX2, NFIX, MEIS2 and TFAP2C have been discussed already (Section 2.3). Then, TCF4 is thought to be important in cortical and hippocampal neurogenesis [48]. STAT3 is thought to be important for neuronal differentiation [49], and more generally the JAK/STAT pathway is known to be important in the transition from neurogenesis to gliogenesis [50]. NFIA is best known as a transcription factor involved in the onset of gliogenesis [51] (i.e., following neurogenesis). Hence, the role of NFIA during neurogenesis is likely to be repressive. The TEAD transcription factors are known to have a role in neural progenitor specification [52], although a specific role for TEAD1 in neurogenesis has not yet been widely reported. Interestingly however, recent work has shown this gene has an important role in cell migration in the aggressive brain cancer glioblastoma [53]. Furthermore, TEAD1 has also been shown to be part of a neuronal transcriptional network which is fundamental to the progression of the pediatric brain cancer medulloblastoma [54]. SOX15 is not yet well known in neural development, although the SOX family of transcription factors are well known as co-factors in lineage specification throughout development [55]. HMBOX1, also known as HOT1, is a gene coding for a protein which binds to telomeres [56]. It’s unclear what its role could be in neurogenesis, although it may be protective as newly generated neurons are known to be hypersensitive to telomere damage [57].

### 2.6 Inferring regulation by NEUROD2

The transcription factor identified in Figure 4 as regulating the largest number of excitatory neuron genes is NEUROD2. Figure 5 shows the local network structure inferred, defined as the genes potentially regulated by NEUROD2. The network structure shown in Figure 5 is obtained from several model fits, each for a different target gene potentially regulated by NEUROD2. This is in contrast to the results for SATB2 in Section 2.3, which is obtained from just one model fit around the target gene SATB2.

**Figure 5:**
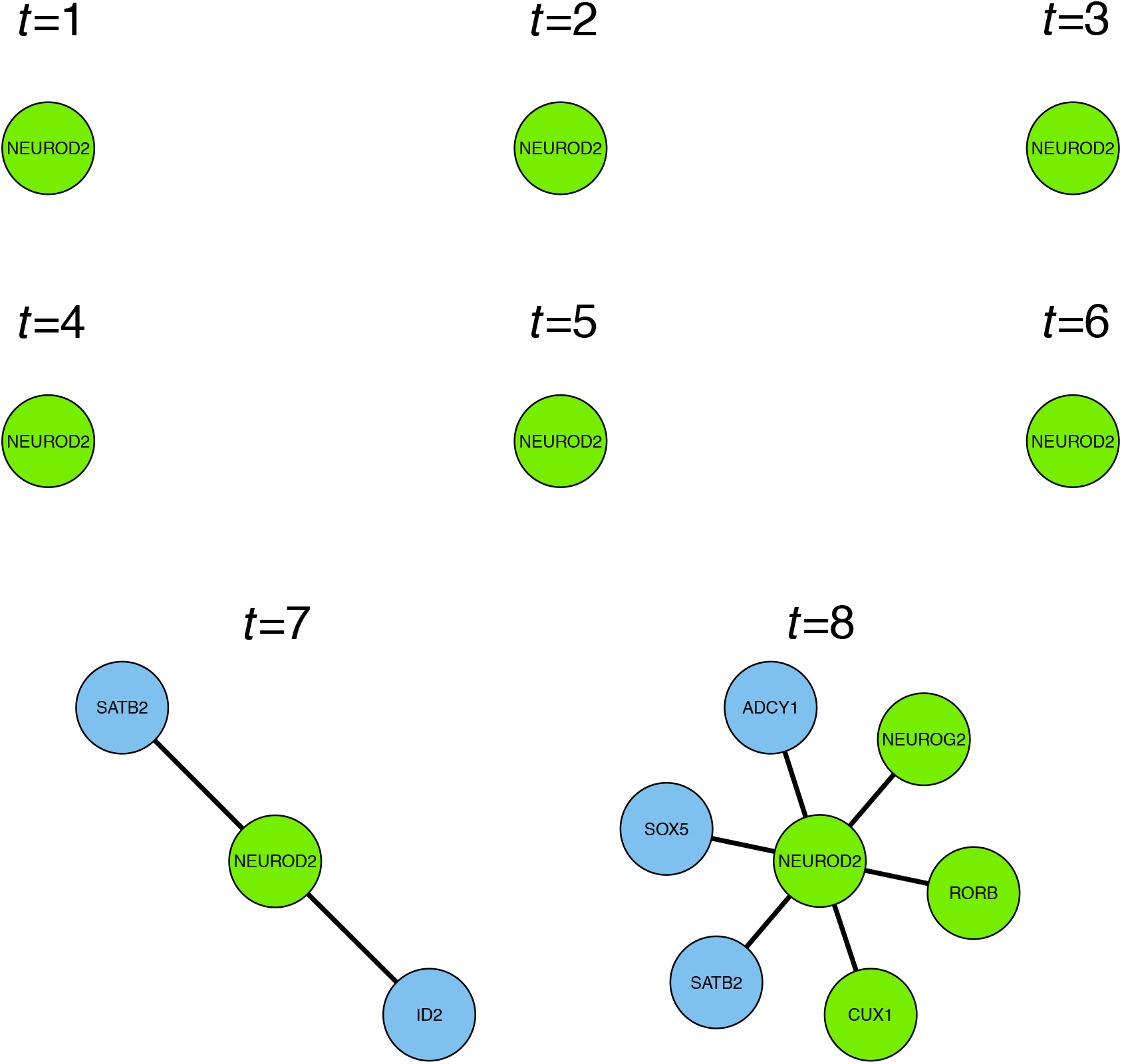
Inferred time-varying genomic regulatory influence of NEUROD2, on the specified set of excitatory neuronal identity genes.

The genes potentially regulated by NEUROD2 (Figure 5) are all of interest to neural development. CUX1, like SATB2, is an important marker of specific neuronal subtypes [4]. NEUROG2 is important for cortical laminar fate specification [58]. RORB is involved in a mutually-repressive interaction with BRN1/2 to specify cortical laminar fate [59]. SOX5 has an important role in neuronal migration and differentiation [60]. ID2 is required for specification of certain types of neuron [61]. Finally, ADCY1 is a neuronal protein thought to have an important role in neuronal signal transduction and synaptic plasticity [62].

### 2.7 Consistent regulation across genes significant in a neuronal subpopulation

Similarly to the results of Section 2.5 (shown in Figure 4), we can also look for consistency of potential transcriptional regulators across target genes taken from a much larger gene-set. To identify such a gene-set, we used LIMMA and edgeR [63, 64] to identify genes which are significantly differentially expressed in a group of cells of interest, when compared to all the other cells in the data-set. We defined this group of cells of interest as those cells assigned to the final time-point in the developmental trajectory. This group of cells is expected to represent a particular excitatory neuronal subtype of interest. Significant genes were defined here as those genes with false discover rate-adjusted *p* < 0.05 in the differential expression analysis, which also increased their expression level in the cells of interest compared to all the other cells. Figure 6 shows the top transcription factors in terms of the numbers of significant genes they are inferred to potentially regulate (the list of genes inferred as regulated by each TF is given in the Appendix).

**Figure 6:**
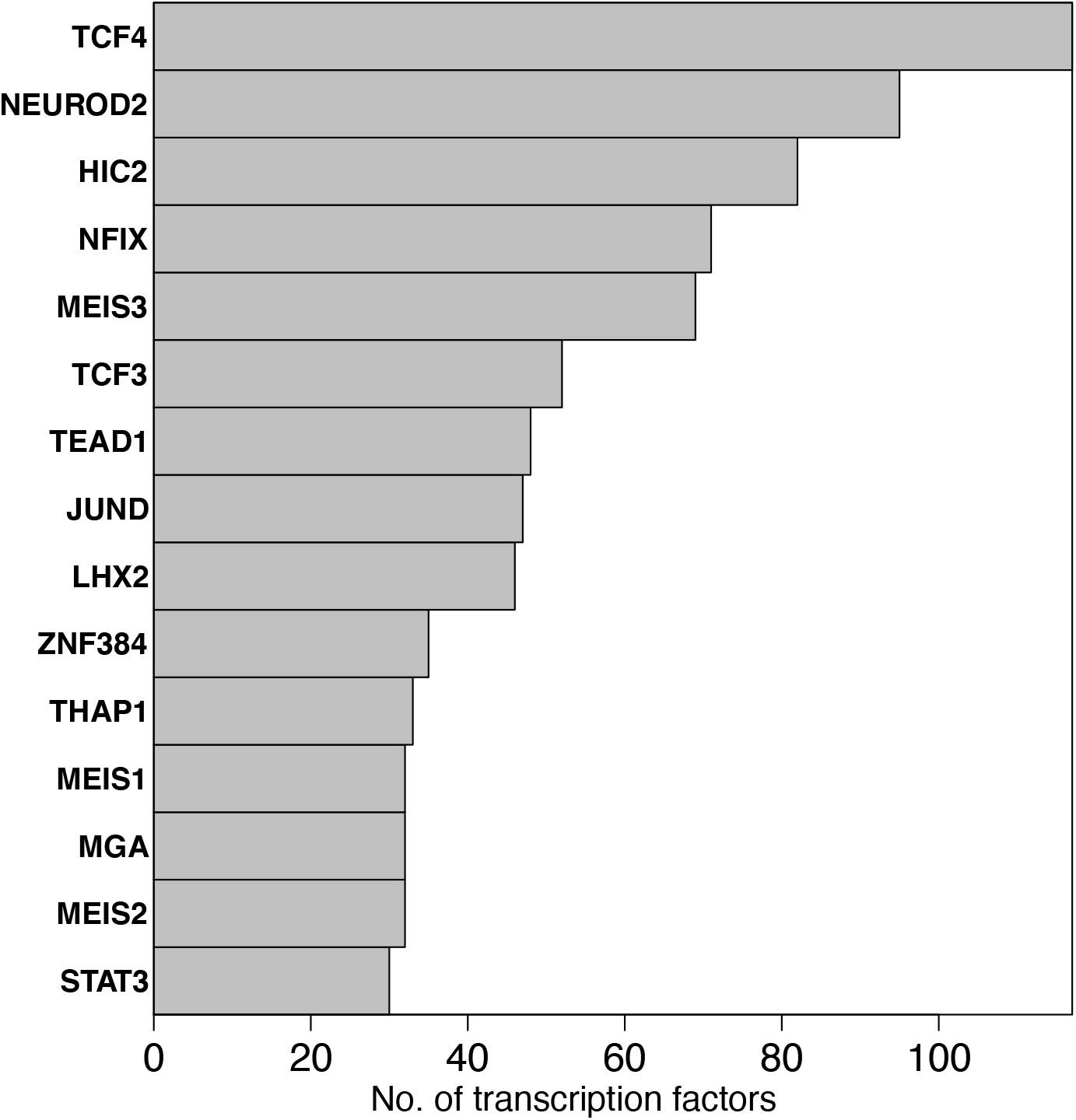
Numbers of neuronal subtype genes of interest, regulated by different transcription-factors.

The transcription factors which appear in Figure 6 are all of interest to neural development. Several have been discussed already, and the rest are now discussed as follows. At the top of the list is TCF4. The role of TCF4 in neural development is not currently well understood, although loss-of-function mutations of this gene have been shown to be responsible for severe neurodevelopmental disorders [65], as well as conferring risk of schizophrenia [66]. However, recent work points to an important role for TCF4 in cortical and hippocampal neurogenesis [48]. Interestingly, HIC2 has no currently reported role in neurogenesis, although parallels have recently been drawn between the role of HIC2 in cardiovascular development, with the role of BRN3A in neural development [67]. MEIS3 is thought to mediate WNT-driven organisation of the neural plate in embryogenesis [68]. TCF3 is known as an inhibitor of neurogenesis, although interestingly this was reported previously in the spinal-cord [69]. JUND is known to have a role in the brain [70], although it may not be involved in development. The zinc-finger gene ZNF384 has previously been reported as having a role in neurogenesis, as part of the gene regulatory circuitry of the ventral neural fate marker PAX6 [71]. THAP1 has previously been reported to have a role in neural development [72]. MEIS1 is known to be an important developmental gene [73]. MGA is ‘MAX gene-associated protein’; MAX is a transcription factor involved with cellular proliferation [36].

### 2.8 Coregulation by NEUROD2 and TCF4

Figure 7 shows a particular co-regulated subnetwork structure of interest, at the final time-point in the inferred dynamic network. This subnetwork structure comprises the neuronal identity genes which are inferred as being potentially regulated by NEUROD2 and TCF4. It is clear from Figure 7 that many of these genes are co-regulated by both these transcription factors.

**Figure 7:**
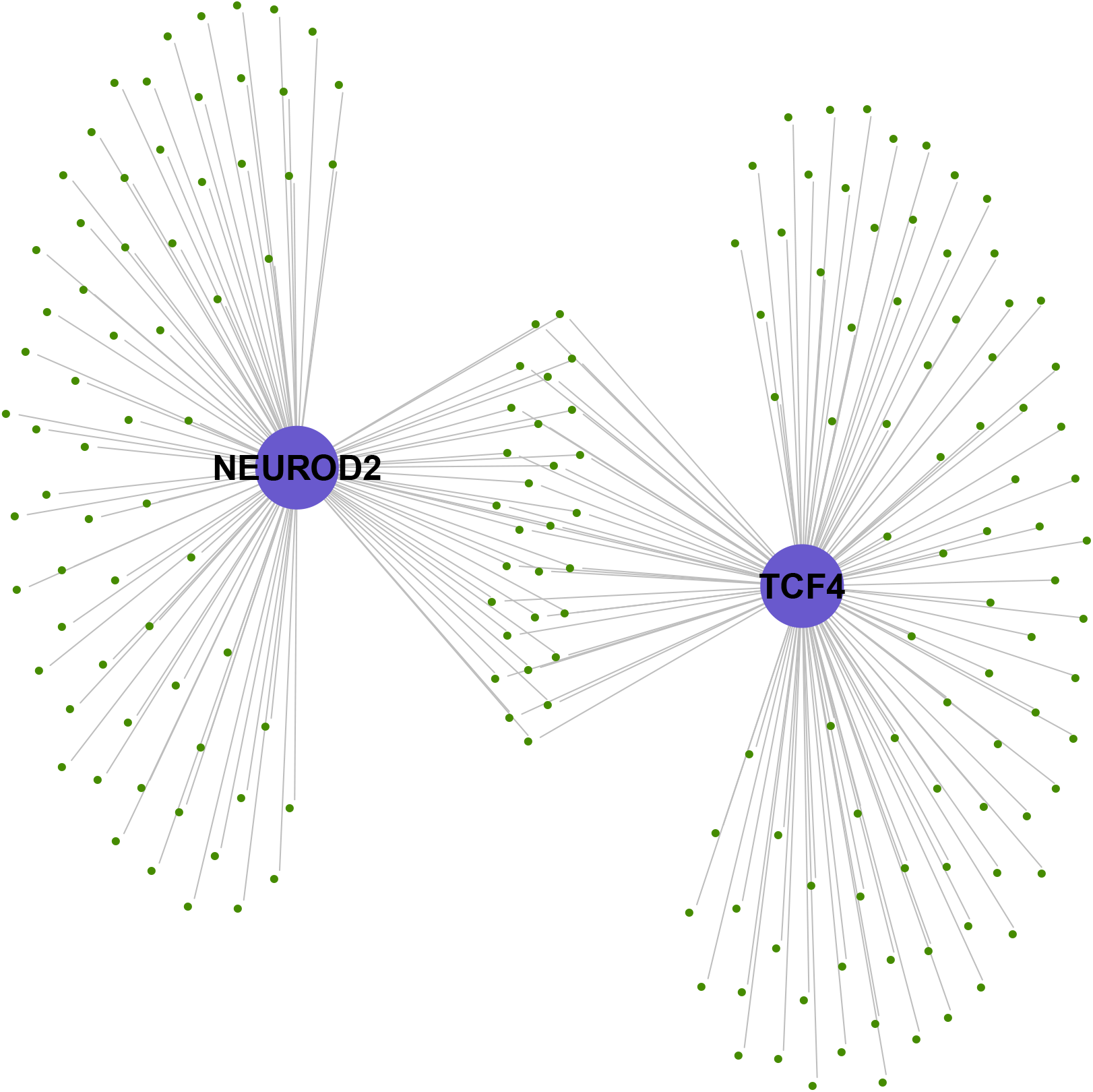
Co-regulation of neuronal subtype genes (shown in green) by the transcription factors TCF4 and NEUROD2 (shown in purple). There are 29 genes connected to both NEUROD2 and TCF4, as well as 66 genes connected to only NEUROD2, and 87 genes connected to only TCF4.

There are 29 genes connected to both NEUROD2 and TCF4 in the subnetwork of figure 7, indicating potential co-regulation by this pair of transcription factors. These 29 genes are: CALCOCO1, CHRDL1, CPSF4, DOCK4, FAM110A, FAM126A, FBLN1, GNG3, HDAC2, HECTD4, IGDCC3, ITPR2, KIAA1324, LRP8, MEX3A, NCS1, NFASC, PGAP1, RBCK1, RBFOX2, RPL37A, SCAF1, SEZ6, SIDT2, SNAP25, TMEM86A, TRIOBP, YWHAG, and ZDHHC20. Several of these are known to be of particular interest, as follows. The gene CHRDL1 promotes neuronal differentiation by inhibiting the important neural development gene BMP4 [74]: the ‘bone morphogenetic proteins’ (BMPs) have a fundamental, but complex, role in cellular specification throughout neural development [75]. The chromatin remodelling gene HDAC2 is ‘histone deacetylase 2’, which has been shown to control the progression of neural precursors to neurons during neural development [76]. The gene KIAA1324 encodes a transmembrane protein, and has been shown to be differentially expressed between the ventricular and subventricular cortical zones [77]. RBFOX2 regulates the alternative splicing of many important neuronal transcripts [78]. DOCK4 is thought to play a fundamental role in formation of neurites and dendrites [79, 80]. NFASC is thought to be involved in neuronal projection morphogenesis [81]. LRP8 is thought to be involved in neuronal migration via its interaction with RELN [82]. GNG3 is expressed in proliferating neural progenitors and immature neurons [83]. IGDCC3 is associated with a committed neuron phenotype [84]. FBLN1 is required for morphogenesis in neural crest-derived structures [85]. SNAP25 is involved in vesicular fusion and neurotransmitter exocytosis, with different isoforms in developing and adult tissue [86].

## 3 Discussion

In this paper, we have presented a new dynamic genomic network model, for inferring patterns of genomic regulatory influence in dynamic cell-biological processes such as development. We have applied this method to genome-wide data from human fetal cortical tissue, finding genomic interactions which are known to be fundamental to excitatory neuron specification. Our method compares very favourably with equivalent findings which we obtain from the same data using a popular method for network inference based on the data correlation matrix.

Our proposed method uses a large repository of publicly-available chromatin accessibility (DNAse-Seq) data, to identify transcription-factor (TF) bindings events that are possible in the neural lineage. It then uses expression data to infer potential regulatory relationships occurring at different times, guided by the possibilities identified in the chromatin accessibility data. We note that a potential regulatory relationship can still be identified if the evidence in the expression data is strong enough, even without corresponding evidence in the chromatin accessibility data. The evidence in the chromatin accessibility data of the possibility of binding of the TF to the regulatory DNA of the target gene effectively reduces the threshold required in the evidence from the expression data, in order to infer a regulatory relationship between these genes.

We have applied our method to data from human tissue, and have interpreted our findings based on knowledge available from the wider literature. Much of what is known about neural development and neurogenesis in mammals is the result of rodent studies. While the neurodevelopmental principles in humans are likely to be similar to rodents in many ways, there must also be key differences due to the much greater size of the human and more generally the primate cortex. Hence, the findings from earlier studies in rodents which we have sometimes cited can only be taken as an indication of genomic regulatory interactions which may take place in human neural development, and particularly cortical neurogenesis.

Recent advances in single-cell genomic profiling include single-cell chromatin accessibility data, including where these measurements are obtained from the same cells as the single-cell RNA-seq measurements. In principle, the methods presented in this manuscript should be directly applicable to such data-sets. However, we note that as single-cell RNA-seq data-sets become larger in size, some trade-off is necessary between run-time and the size of the data-set.

Our method is computationally efficient, and can be applied to genome-wide data with tens of thousands of transcripts. However, we note that in order to define and describe genomic interactions which are more specifically mechanistic, a finer-grained model will be needed. This finer-grained model will necessarily be more complex, and thus would not be feasible to run at this genome-wide scale. Hence, the method we propose is most appropriate for a course-grained genome-wide discovery or exploration stage. This discovery stage can then be followed by the finer-grained stage of mechanistic modelling, which should also incorporate experimental validation.

## 4 Conclusions

The method we propose here provides a new mathematical and computational tool, which could be used together with targeted experiments in order to reveal important new functional genomic regulatory processes in mammalian development.

## Declarations

### Ethics approval and consent to participate

Not applicable.

### Consent for publication

Not applicable.

### Availability of data and materials

The data-sets analysed during the current study are available in the NCBI database of genotypes and phenotypes (*dbGaP*) under accession number *phs000989.v3* with URL https://www.ncbi.nlm.nih.gov/projects/gap/cgi-bin/study.cgi?study_id=phs000989.v3.p1 and from *ENCODE* under accession numbers *ENCFF196CDZ.bam ENCFF436NOR.bam ENCFF807SCV.bam ENCFF808BII.bam ENCFF537DKA.bam ENCFF108NTA.bam ENCFF431RLT.bam ENCFF542AQF.bam ENCFF542NTO.bam ENCFF602FMG.bam ENCFF846NFV.bam ENCFF848SVJ.bam ENCFF948DOQ.bam* and *ENCFF952QLZ.bam* with URLs https://www.encodeproject.org/files/ENCFF196CDZ/ https://www.encodeproject.org/files/ENCFF436NOR/ https://www.encodeproject.org/files/ENCFF807SCV/ https://www.encodeproject.org/files/ENCFF808BII/ https://www.encodeproject.org/files/ENCFF537DKA/ https://www.encodeproject.org/files/ENCFF108NTA/ https://www.encodeproject.org/files/ENCFF431RLT/ https://www.encodeproject.org/files/ENCFF542AQF/ https://www.encodeproject.org/files/ENCFF542NTO/ https://www.encodeproject.org/files/ENCFF602FMG/ https://www.encodeproject.org/files/ENCFF846NFV/ https://www.encodeproject.org/files/ENCFF848SVJ/ https://www.encodeproject.org/files/ENCFF948DOQ/ and https://www.encodeproject.org/files/ENCFF952QLZ/

### Author contributions

TB conceived and designed the study, carried out all analyses, and wrote the manuscript.

## Acknowledgements

Not applicable.

